# Impact of porcine cytomegalovirus on long-term orthotopic cardiac xenotransplant survival

**DOI:** 10.1101/2020.04.07.029702

**Authors:** Joachim Denner, Matthias Längin, Bruno Reichart, Luise Krüger, Uwe Fiebig, Maren Mokelke, Julia Radan, Tanja Mayr, Anastasia Milusev, Fabian Luther, Nicoletta Sorvillo, Robert Rieben, Paolo Brenner, Christoph Walz, Eckhard Wolf, Berit Roshani, Christiane Stahl-Hennig, Jan-Michael Abicht

## Abstract

Xenotransplantation using pig organs has achieved survival times of more than 195 days in pig orthotopic heart transplantation into baboons. Here we demonstrate that in addition to an improved immunosuppressive regimen, non-ischaemic preservation with continuous perfusion and control of post-transplantation growth of the transplant, prevention of transmission of the porcine cytomegalovirus (PCMV) plays an important role in achieving long survival times. For the first time we demonstrate that PCMV transmission in orthotopic pig heart xenotransplantation was associated with a reduced survival time of the transplant and increased levels of IL-6 and TNFα were found in the transplanted baboon. Furthermore, high levels of tPA-PAI-1 complexes were found, suggesting a complete loss of the pro-fibrinolytic properties of the endothelial cells. These data show that PCMV has an important impact on transplant survival and call for elimination of PCMV from donor pigs.

---

Recently, consistent success in life-supporting (orthotopic) porcine cardiac xenotransplantation has been reported^1^. In that study hearts from α1,3-galactosyltransferase-knockout (GTKO) pigs that express human membrane cofactor protein (CD46) and human thrombomodulin (hTM) had been transplanted into baboons and survival times up to 195 days were achieved. This is a milestone on the way to clinical cardiac xenotransplantation which is urgently needed: The supply of human organs does not match the needs and many patients with terminal cardiac failure die while being on the waiting list.

Xenotransplantation with genetically modified porcine organs, as an alternative to allogenic (human-to-human) procedures may be associated with the transmission of porcine microorganisms, among them the porcine endogenous retroviruses (PERVs). PERVs are integrated in the genome of all pigs and they are able to infect human cells^2^. However, until now no PERV transmission was observed in the first preclinical (for review see ref. 2) and clinical xenotransplantation trials^3,4^. Whereas PERV-A and PERV-B, which are present in all pigs, infect human cells, PERV-C infects only pig cells and is not present in all pigs^2^. However, recombinations between PERV-A and PERV-C can occur and the recombinant PERV-A/C is characterised by a higher replication competence^5,6^. Therefore, it is highly recommended to use PERV-C-free pigs for xenotransplantation. Interestingly, using CRISPR/Cas all retroviral sequences can be inactivated in the pig genome^7,8^, however, it is still unclear whether this is needed for a safe xenotransplantation^9^.

From the other viruses widely distributed in pigs, the porcine cytomegalovirus (PCMV) is of great concern^10^. PCMV is related with the human cytomegalovirus (HCMV), also called human herpesvirus 5 (HHV-5). HCMV causes fatal infections in human organ transplant recipients if not treated, leading to end-organ disease, such as gastrointestinal ulceration, hepatitis, pneumonitis or retinitis. HCMV can also lead to systemic infection and disease once a threshold value of virus load is exceeded^11^. In fact, HCMV has also been detected in the bowel mucosa where its reactivation has been suggested to lead to IL-6 release and inflammatory bowel disease^12^.

Meanwhile it was shown that PCMV is a roseolovirus and more closely related with human herpesviruses 6A, 6B and 7 (HHV-6A, HHV-6B, and HHV-7)^13^. Therefore the thermology PCMV is to a certain degree misleading and should actually be porcine roseolovirus (PCMV/PRV)^14^. The International Committee on Taxonomy of Viruses (ICTV) classified this virus as suid betaherpesvirus 2^15^. The closely related HHV-6 and HHV-7 were reported to be associated with numerous diseases, e.g., liver failure^16^, multiple sclerosis^17^ and Alzheimer disease^18^. Furthermore, HHV-6 was found to promote cancer development^19^ and accelerate acquired immunodeficiency syndrome (AIDS) in humans and monkeys^20,21^, possibly by its immunosuppressive property^21-25^.

Recently, by analysing a baboon recipient of orthotopic pig heart transplantation with a relatively short survival time (29 days) and hepatic failure, PCMV/PRV infection was observed in the recipient^26^. In addition, PCMV/PRV transmission was also found in two other baboon recipients with 4 and 40 days of transplant survival. Immunohistochemical studies of the recipient baboons showed PCMV/PRV-expressing cells in all organs of the animal, most likely representing disseminated pig cells^27^. These data, together with similar data on pig kidney xenotransplantions in non-human primates^28,29^, suggest that PCMV/PRV significantly reduces the survival of pig xenotransplants. However, the mechanism through which PCMV/PRV reduces transplant survival is still unclear^30^.

To better understand the impact of PCMV/PRV on pig transplant survival in orthotopic heart transplantation, numerous donor pig - baboon recipient pairs were retrospectively analysed for PCMV/PRV transmission. Here, we demonstrate for the first time that PCMV transmission in orthotopic pig heart xenotransplantation is associated with a reduced survival time of the transplant and indicate an impact of PCMV/PVR on cytokine release and coagulation. Furthermore, we show that other porcine viruses, which could potentially impact xenotransplant survival, including PERVs, hepatitis E virus (HEV), three porcine lymphotropic herpesviruses (PLHV) and the porcine circoviruses (PCV) 1 and 2, had not been transmitted.

## Results

### Transmission of PCMV/PRV into baboons after orthotopic pig heart transplantation

All baboon recipients received an immunosuppression including an induction therapy with an anti-CD20 antibody, anti-thymocyte-globulin and a monkey-specific anti-CD40 monoclonal antibody or humanized anti-CD40PASylated as described in detail^1^. Three groups of animals have been transplanted. In group I, donor organs were preserved with two clinically approved crystalloid solutions, the animals survived for less than 30 days (animal C, Table 1) and suffered from perioperative cardiac xenograft dysfunction (PCXD). To reduce the PCXD, in group II the pig hearts were preserved with an oxygenated albumin-containing hyperoncotic cardioplegic solution. Three of the four animals of this group lived for 18 (animal F), 27 (animal H) and 40 (animal I) days. A diastolic heart failure and subsequent congestive liver damage resulting from massive cardiac overgrow were observed in these animals. To prevent this, in group III baboon recipients were weaned from cortisone at an early stage and received antihypertensive treatment since pigs have a lower systolic blood pressure than baboons. In addition, a temsirolimus medication was used to counteract cardiac overgrowth. Two recipients in this group survived for 195 (animal O, Table 2) and 182 (animal N) days^1^.

**Table 1.**
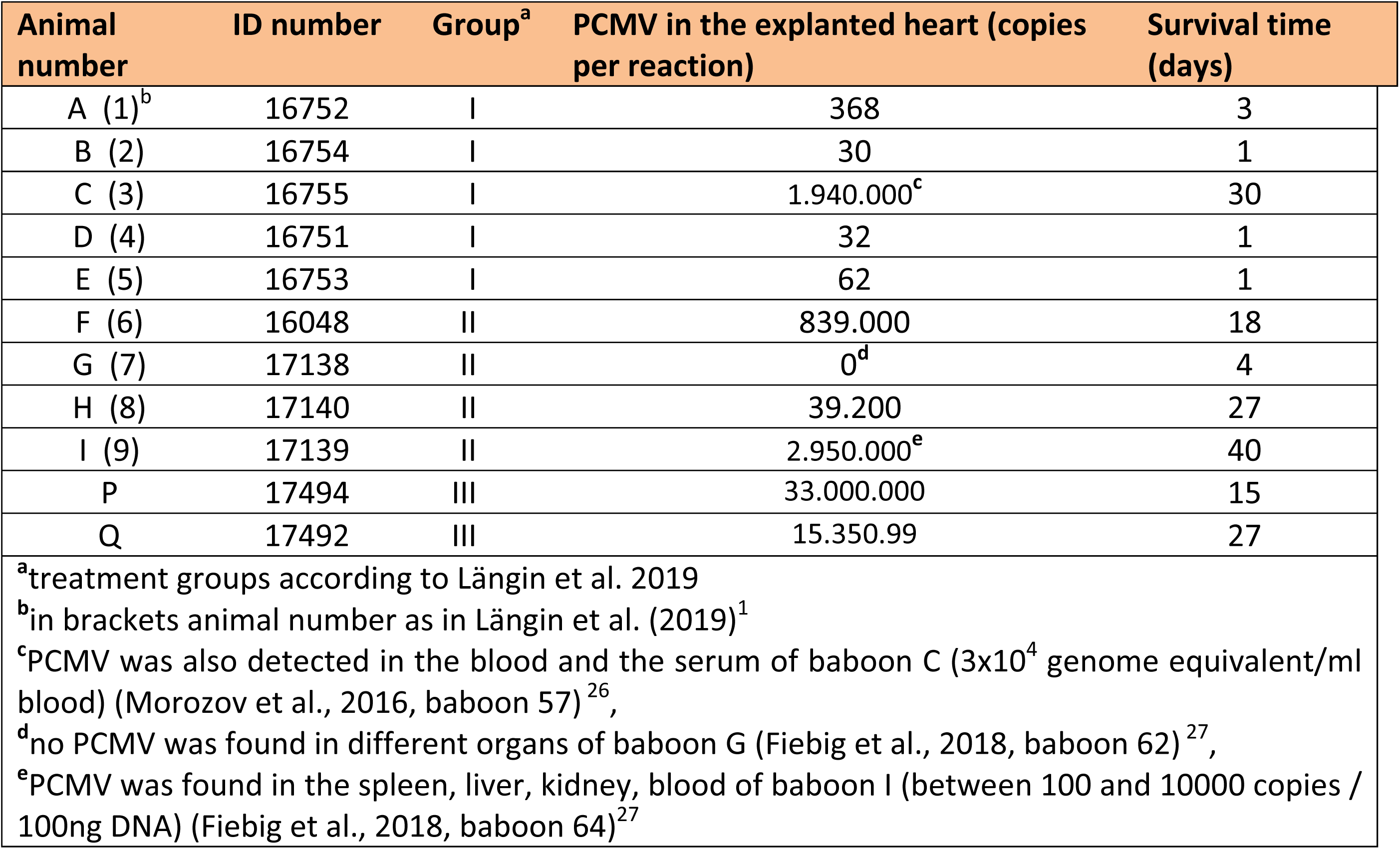
PCMV infection in explanted pig hearts from different baboons.

**Table 2.**
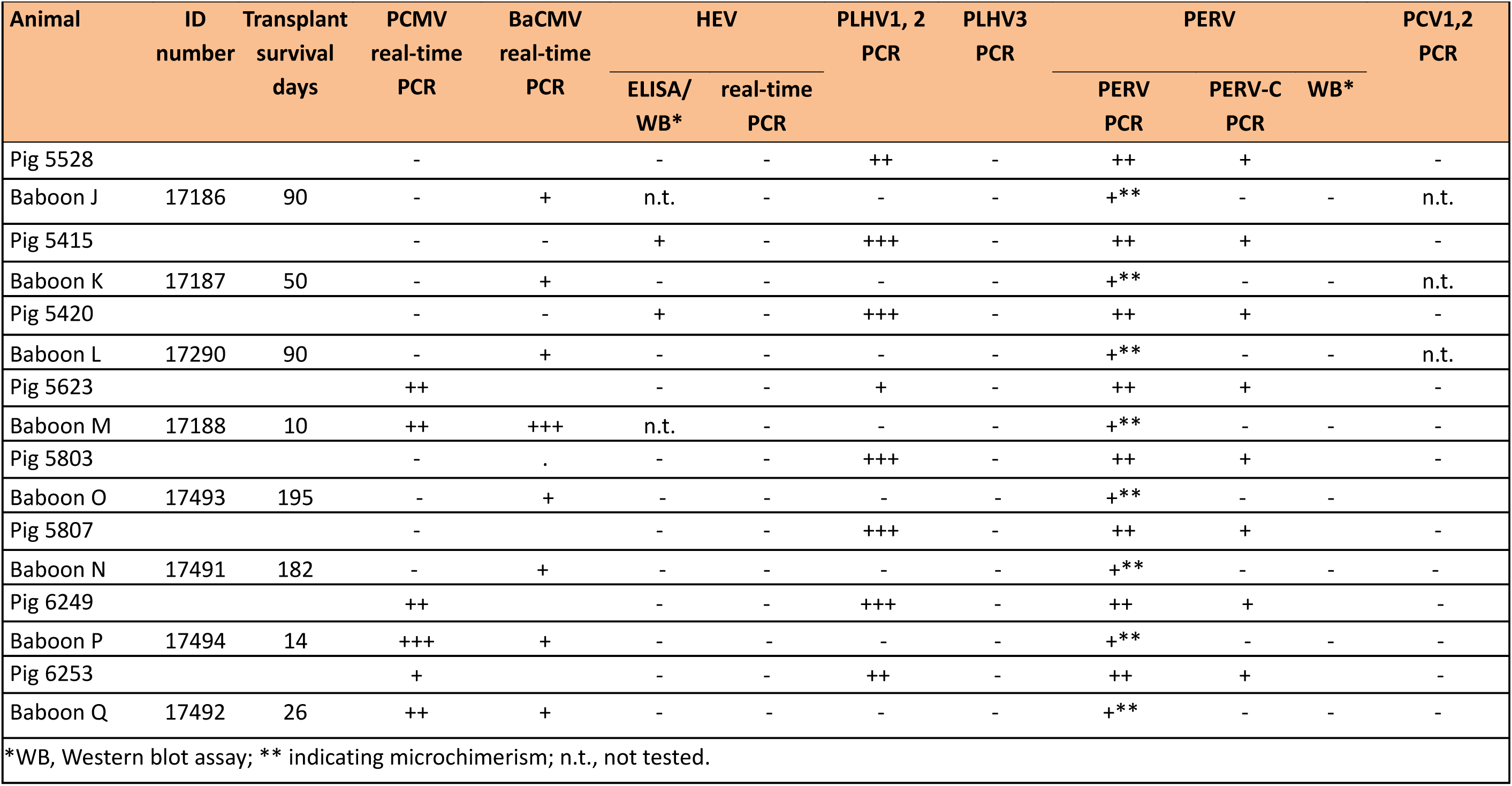
Testing donor pigs and recipient baboons of transplantation group III for different viruses.

When the recipient baboons organs from group I and group II were analysed for PCMV/PRV transmission, PCMV/PRV was detected not only in the explanted heart but also in several different organs from all baboons except one (G; Table 1). Some of these animals (B, D, E) survived only one day and showed very low virus load in the explanted heart. However, after only three days an increase of the virus load was observed (animal A). When analysing the animals with the longer survival time, e.g., animal F with 18 days, animal H with 27 days, animal C with 30 days and animal I with 40 days, it became evident that PCMV/PRV replicates over time. Animal I (number 64 in publication 27) had not only a very high virus load in the explanted heart, but a high virus load was also found in the spleen, kidney, and blood (between 100 and 1000 copies / 100ng DNA) and cells expressing PCMV/PRV proteins were found in all organs of the transplanted animal^27^. The longer the transplant was in the recipient the higher was the virus load in the explanted and formalin/ethanol fixed pig heart (Table 1). This rule is operative only to a certain threshold of the virus load, above this threshold the survival time decreases due to the pathogenic effect of the virus.

When the recipient baboons of group III (early cortisone tapering, antihypertensive treatment and temsirolimus) and their donor pigs were analysed, PCMV/PRV was found to be absent in the donor pigs and consequently in the transplanted baboons of animals with the longest survival times such as baboon J (90 days), baboon K (50 days), baboon L (90 days), baboon N (182 days) and baboon O (195 days) (Table 2). In contrast, PCMV/PRV was found in the donor pigs of transplanted baboons that showed a shorter survival rate like baboon P (14 days), baboon Q (26 days) and M (euthanized 10 days after transplantation because of severe iatrogenic complication) (Table 2). In the hearts explanted from animals P and Q astonishingly high copy numbers of PCMV/PRV were detected. Since all these data were obtained testing the formalin/ethanol fixed pig hearts, in addition the virus load in the frozen right ventricle of the heart explanted from baboon P was analysed and 48 million copies were found (33 million in the fixed heart). This is in the same magnitude, indicating that testing of fixed and frozen materials give approximately the same result.

### Distribution of PCMV/PRV in pig and baboon organs

Using real-time PCR, the copy number (viral load) of PCMV/PRV was analysed in PBMCs and different organs of all donor pigs and in the organs of the transplanted baboons after removal of the transplanted heart, which was also analysed. As an example we illustrate in Figure 1 the pair donor pig 6249 and the recipient baboon P. All other animals analysed had a similar pattern. With exception of the skin, the viral load was high in all baboon organs (spleen, liver, muscle, kidney lung and lymph nodes) and donor pig organs (lung, spleen, liver, kidney and PBMCs), in agreement with previous results^27^. The highest viral load was found in the explanted pig heart (Figure 1). This is the main place of virus replication and this replication is possible due to the absence of the pig immune system, which could have reacted against the virus in the donor pig, and due to the strong immunosuppression in the baboon inhibiting its immune system.

**Figure 1.**
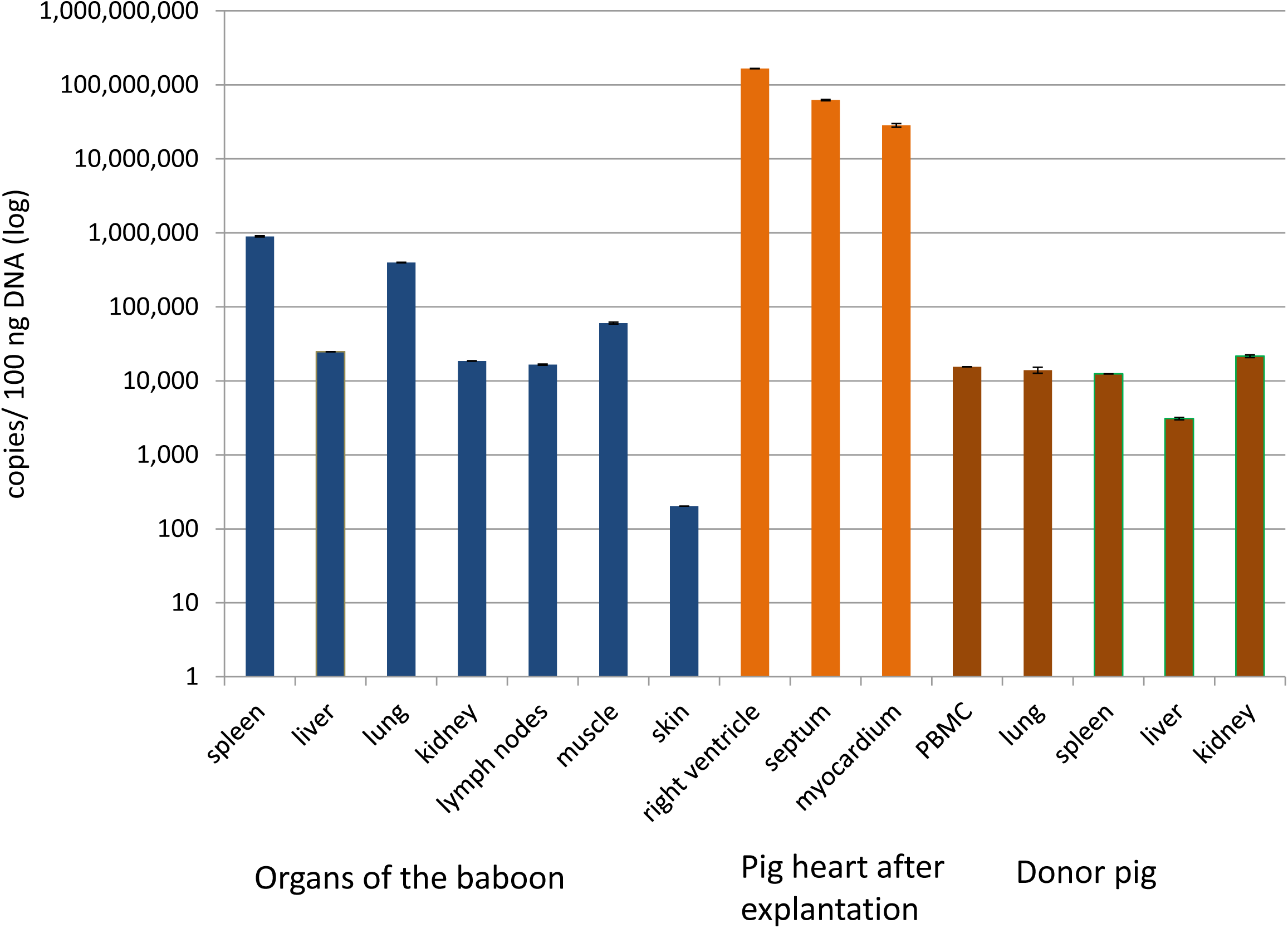
Viral load (copies per 100ng DNA) in different organs of baboon P after explantation of the pig heart, in PBMCs and different organs of the donor pig and in the pig heart after explantation as measured by real-time PCR.

### Pathological findings and modulation of cytokine release

As mentioned above, baboons N and O from treatment group III, showed a longer survival rate, 182 and 195 days respectively, compared to baboons P73 and Q, which survived only 15 and 27 days. Interestingly, both baboons P and Q were characterized by pathological and inflammatory changes, some of which might be the result of a PCMV/PRV infection, bearing in mind the broad disease spectrum induced by herpesviruses. Both animals showed signs of low-output heart failure at the end of the experiments, resulting in multi organ dysfunction as indicated by the pathological increase in functional parameters of liver, pancreas and kidney such as aspartate aminotransferase, creatine kinase, and lactate dehydrogenase. Decreased abdominal perfusion probably caused loss of mucosal barrier function in the gut, leading to translocation of intestinal microbes and to a strong increase of levels of interleukin-6 (IL-6) (Figure 2). In fact, Klebsiella pneumonia was found in blood cultures of baboon Q one day before euthanasia. The increase in IL-6 most probably is not due to treatment with an IL-6-receptor antagonist, since treatment began at the very beginning of transplantation and was also given to animals N and O that did not show high plasma levels of the cytokine. To confirm these data and to perform a broader analysis of further pro- and anti-inflammatory cytokines, a cytometric bead array (CBA) assay for non-human primate Th1/Th2 cytokines as well as an IL-10 ELISA were performed. The obtained data confirmed the strong increase of IL-6 (Figure 3A), and also revealed a substantial increase of tumour necrosis factor (TNF) in baboons P73 and Q74 with the PCMV/PRV-positive hearts (Figure 3B). No alterations were observed in serum levels of the proinflammatory cytokines interferon γ (IFN γ) (Figure 3F) and interleukin 2 (IL-2) (Figure 3D) as well as the anti-inflammatory cytokines interleukin-4 (IL-4) (Figure 3E), interleukin-5 (IL-5) (Figure 3F), and interleukin 10 (IL-10) (data not shown).

**Figure 2.**
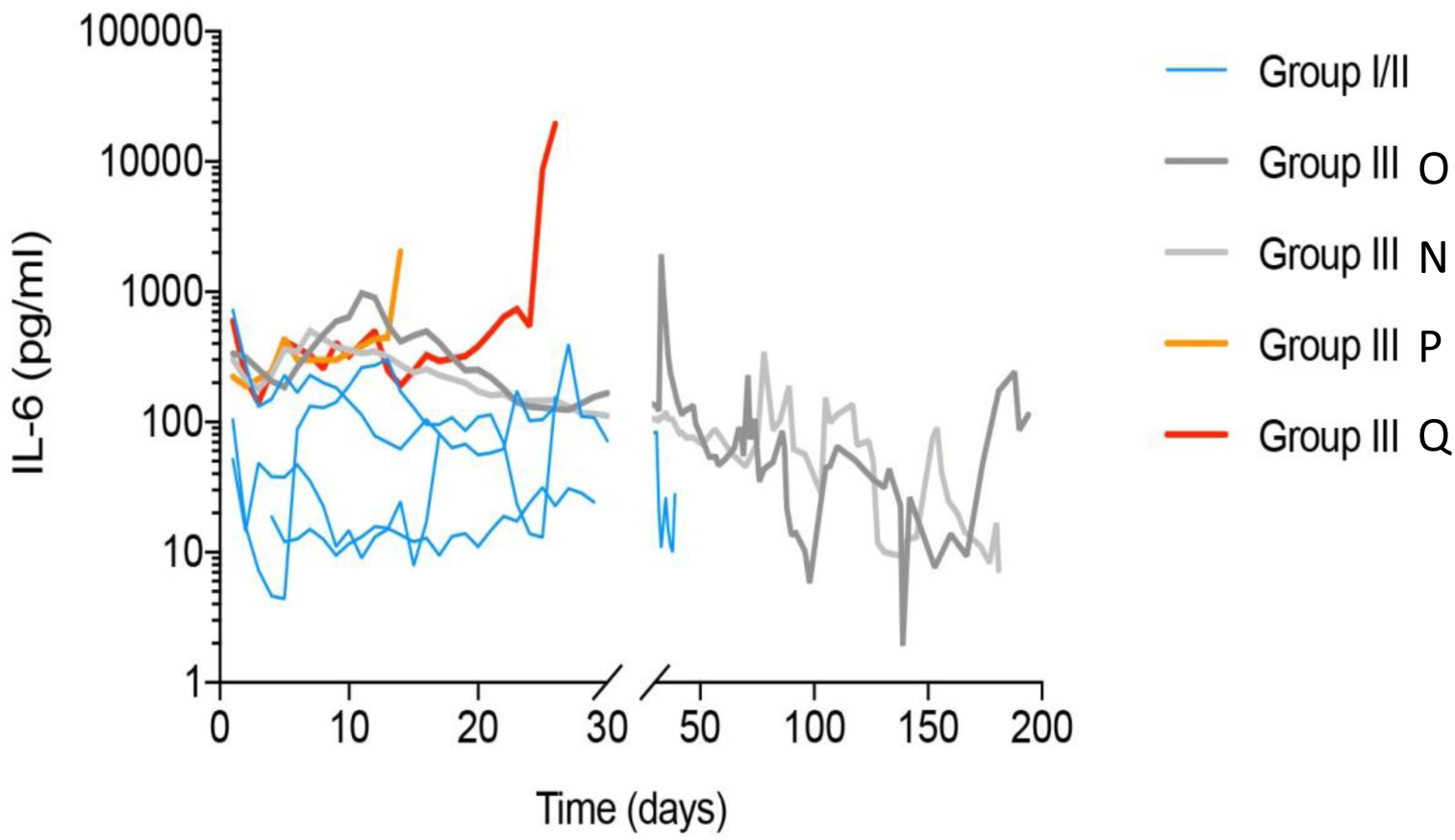
Kinetics of the IL-6 amount in the blood of transplanted baboons as measured by ELISA. The baboon recipients of the treatment group III are indicated (N, O, P, Q).

**Figure 3.**
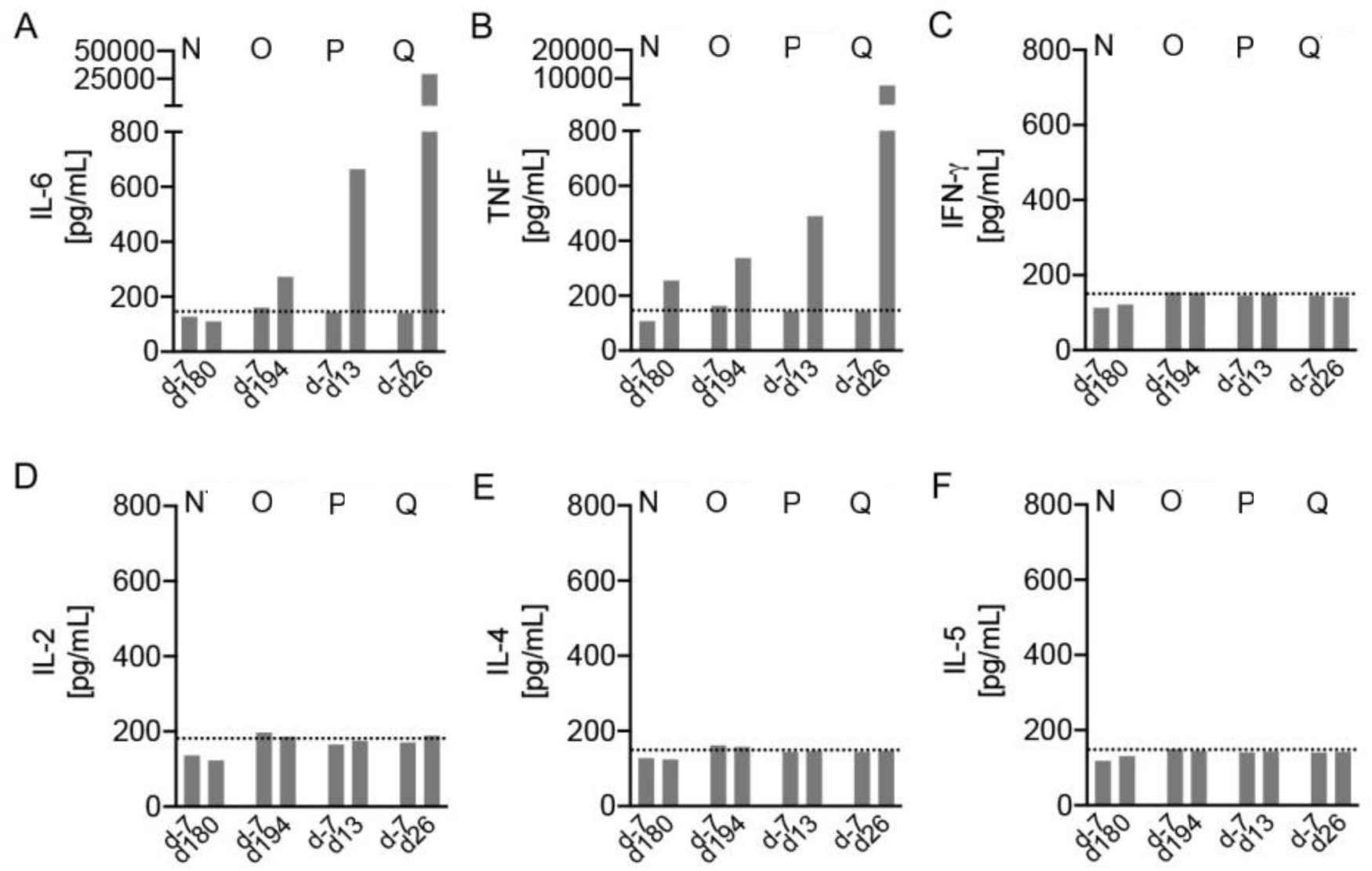
Increased IL-6 and TNF serum levels in baboons with PCMV/PRV-positive hearts. BD CBA Non-human primate Th1/Th2 kit was used to measure plasma levels of (A) IL-6, (B) TNF, (C) IFN γ, (D) IL-2, (E) IL-4 and (F) IL-5 one week before transplantation (day -7) and at the end of experiment (day 13, 26, 180 and 194, respectively) in baboons N, O, P and Q. Dotted lines indicate lowest standard concentration.

Interestingly, histological examination of the explanted pig hearts from baboons N, O, P, and Q revealed marked perivascular and moderate edema in otherwise unremarkable myocardial tissue (Figure 4). In particular, no morphologic signs of cellular or antibody-mediated transplant rejection was found, and in parallel no elevated levels of non-galactose-a1,3-galactose reactive IgM and IgG were measured (not shown), indicating that immunological rejection was not the cause of the reduced survival observed in animals P and Q.

**Figure 4.**
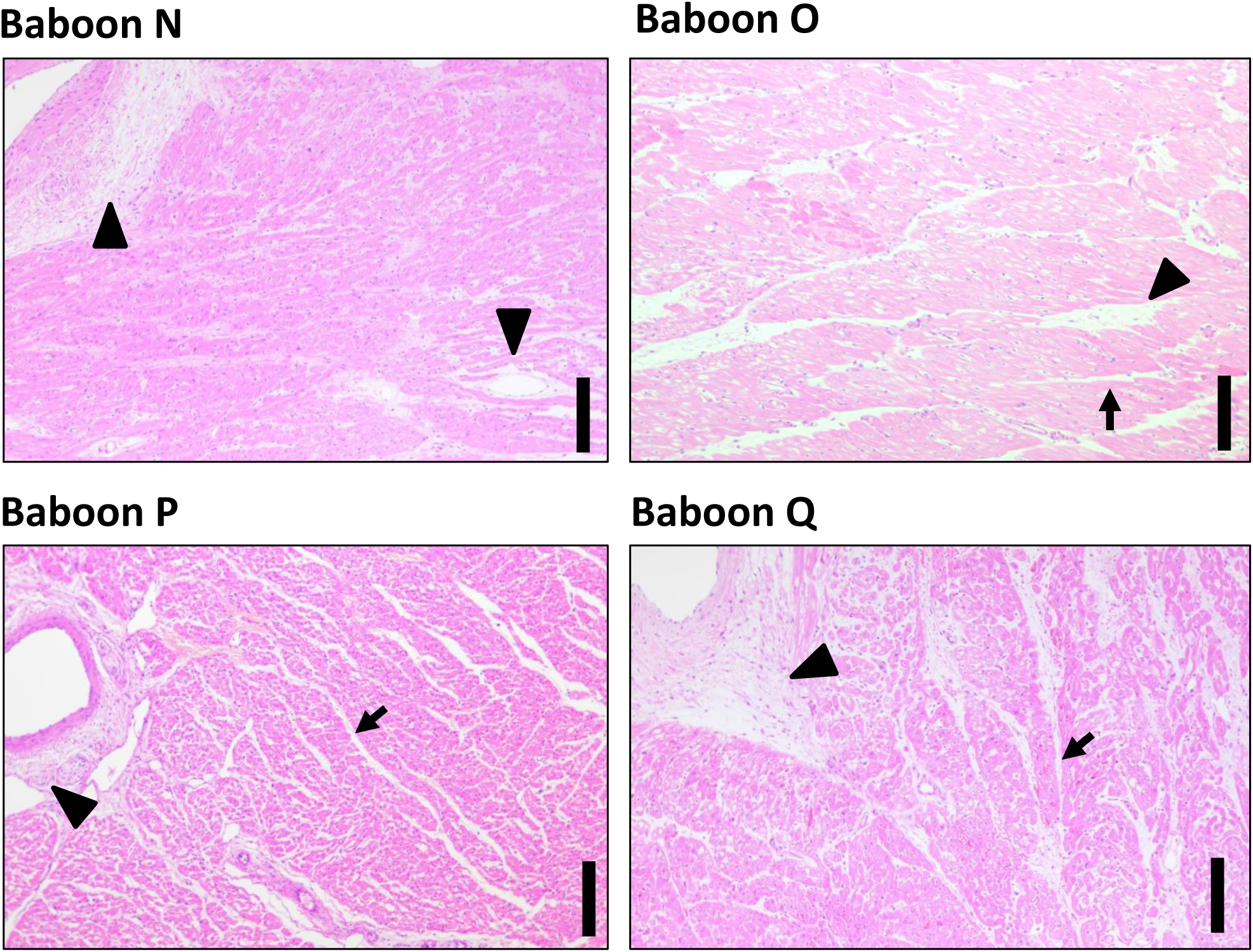
Histopathologic evaluation of the explanted pig hearts from baboons N to Q. Arrowheads mark perivascular and arrows moderate edema in otherwise unremarkable myocardial tissue in all specimens. Scale bars indicate 250μm

### High levels of tPA-PAI-1 complex

To analyse whether animals infected with PCMV/PRV could present alteration in the coagulation system, tissue plasminogen activator (tPA) and plasminogen activator inhibitor 1 complexes (tPA-PAI-1) were measured in plasma samples by an in-house developed Bio-Plex assay^31^. Interestingly, we observed that baboons P and Q, which presented clear pathological alterations, also had very high levels of tPA-PAI-1 complexes (Figure 5). This indicates a hypercoagulable state of the two animals and a decrease in fibrinolysis, originated most probably by endothelial cell damage.

**Figure 5.**
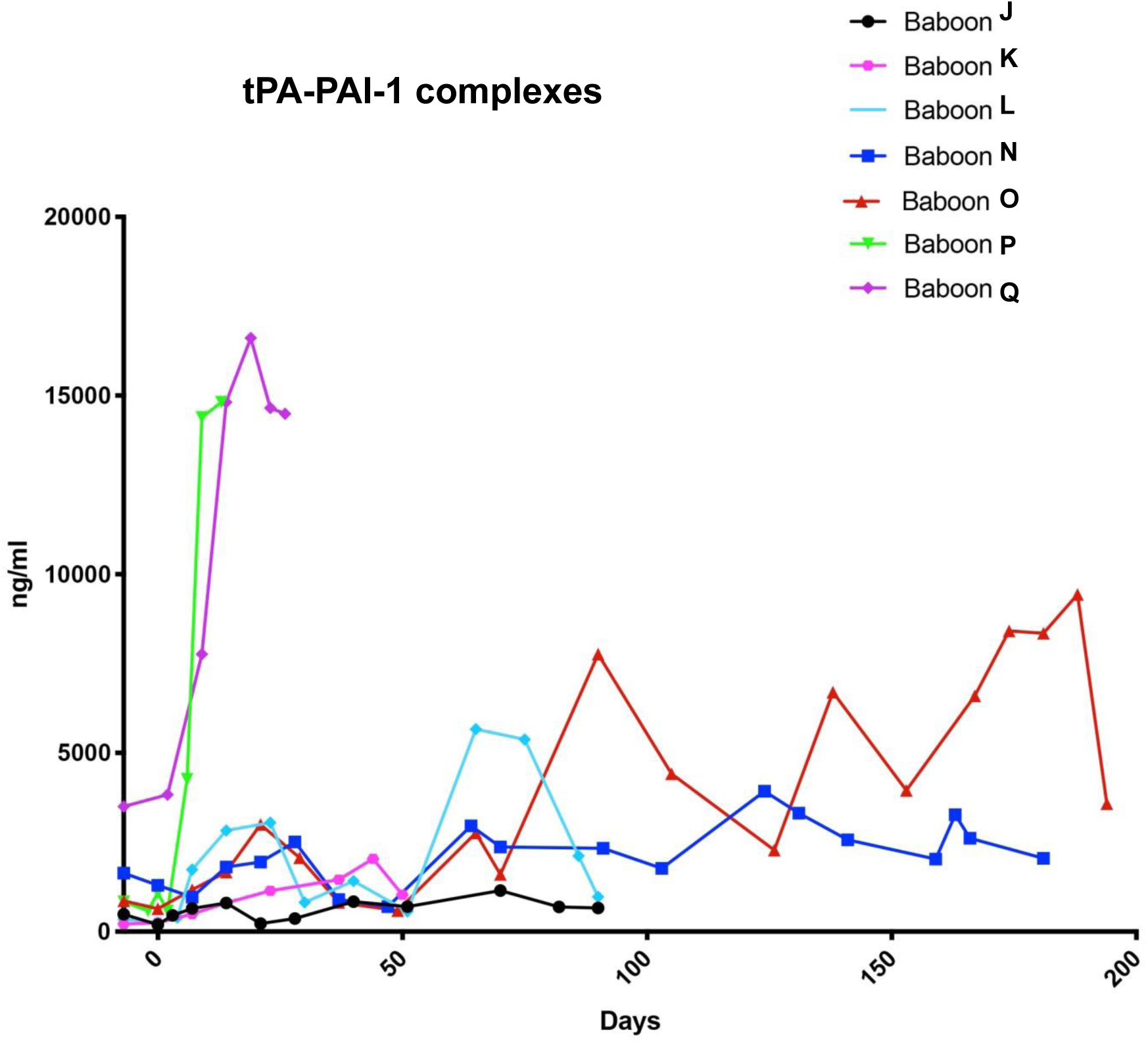
Measurement of tPA-PAI-1 complexes in the blood of baboons J65-Q74.

### Expression of BaCMV after PCMV/PRV transmission

Since all baboon recipients carried BaCMV, we looked into a possible activation of this virus, which may also influence the survival time of the transplant and recipient. Recently a significantly increased expression of BaCMV in plasma of baboon I after transplantation of a PCMV/PRV-positive pig heart was measured^27^. Although, analysis of BaCMV in all baboon organs (liver, kidney, spleen and lung) showed a high virus load a clear and measurable increase of the BaCMV virus load in the blood after transplantation as was found in animal I was not observed (Table 2). Moreover, no differences in the BaCMV viral load in baboons with a long or a short survival time was observed. This highly suggests that activation of BaCMV is not the cause of early loss of the transplanted pig heart.

### Prevalence and transmission of other porcine viruses

In order to analyse whether other porcine viruses were present in the donor pig and were transmitted to the recipient, which also may influence the survival time of the xenotransplant, both Western blot and RT-PCR analysis was performed to detect PERV, PLHV, PCV and HEV in both pigs and baboon tissues. PLHV-1 and PLHV-2 had been detected in all donor pigs used in this study. Despite this, no transmission of these viruses to the recipient baboons was observed (Table 2). Although Western blot analyses showed HEV infection in the donor pigs of baboons G, I, K and L, detection of HEV by RT-PCR gave negative results in all donor pigs, indicating that the virus was not present in their blood. However, transmission to the recipients was not observed (Table 2). Indeed, all baboons were negative for HEV infection independently to the test used for analysis, e.g., no positive RT-PCR reactions and no positive ELISA or Western blot analyses were found.

PERV was detected as expected in all donor pigs by PCR using specific primers binding to a highly conserved region in the polymerase (pol) gene (Table 2). In fact, these primers recognise both PERV-A, PERV-B and PERV-C. Since PERV-A and PERV-B is present in all pigs^2^, a PCR using primers specific for the env region of PERV-C was performed, and PERV-C was detected in all donor pigs.

PERV was also analysed in baboon samples using the pol specific PCR, detecting PERV-A, PERV-B and PERV-C. As shown in Table 2, these sequences were detected in all baboon blood samples, however the detection of PERV in the blood samples might be due to cellular DNA from circulating pig cells, a phenomenon called microchimerism. This was confirmed by the finding of porcine glycerinaldehyde-3-phosphat-dehydrogenase (GAPDH) by a specific PCR. Since the copy number of PERV-C in pig cells is much lower compared to the copy number of PERV-A and PERV-B, PERV-C sequences could not be detected in the blood of the baboons. Moreover, Western blot analysis showed no anti-PERV antibodies in all baboon samples, clearly indicating absence of infection (Supplementary Figure 1). In summary, no other pig viruses which could influence the survival time of the transplant had been transmitted to the recipient baboons.

## Discussion

Recently, survival times of 182 and 195 days after preclinical orthotopic pig-to-baboon xenotransplantation were achieved by several measures that include an improved immunosuppressive regimen, non-ischaemic preservation with continuous perfusion and control of post-transplantation growth of the transplant^1^. Here, for the first time, we show that the use of organs from PCMV/PRV-positive pigs in this life-supporting model is associated with a reduction of the transplant survival time.

Our data on reduction of the survival time of PCMV/PRV-positive pig hearts orthotopically transplanted into baboons supports previous findings in other preclinical trials on kidney xenotransplantations^28, 29, 32-37^. In these studies, PCMV/PRV was found in all organs of the baboon recipients, with the highest copy numbers in liver, lung and kidney^33^. An increased PCMV/PRV virus load was found in 22 different pig xenotransplants with a survival time between 7 and 32 days in baboons^33^. The presence of PCMV/PRV was usually found associated with consumptive coagulopathy (CC)^34^. However, there was no correlation between PCMV/PRV infection and CC, but lower levels of PCMV/PRV infection were always associated with a prolonged transplant survival^34^. When a heterotopic pig heart transplantation was performed in baboons, the survival time of the transplant was shorter when the organs were PCMV/PRV-positive (median 20 days), compared with PCMV/PRV-negative transplants (median 53 days)^35^. Most importantly, early weaning was shown to prevent PCMV/PRV infection of the pigs^35^. Organs from PCMV/PRV-free animals due to early weaning did not induce CC. Unfortunately, PCMV/PRV had been shown to have a reduced susceptibility to ganciclovir which is very effective in inhibiting HCMV^36^. Ganciclovir was also given to the baboons in the present study. When kidneys from GTKO pigs were transplanted into baboons, the median survival time of the kidneys from PCMV/PRV-positive animals was 14.1 days, the survival time of kidneys from PCMV/PRV-negative animals was 48.8 days and that of kidneys from PCMV/PRV-negative animals after a Caesarean delivery was 53 days^28^, clearly demonstrating that the transmission of PCMV/PRV reduced significantly the kidney transplant survival. A similar effect was observed, when GTKO pig kidneys were transplanted into cynomolgus monkeys. Whereas the survival time of kidneys from PCMV/PRV-free animals was 28.7 days, the presence of PCMV/PRV reduced the survival time to 9.2 days^29^.

Our findings that PCMV/PRV induces IL-6 and TNF release and increases the amount of t PAI-1/tPA complexes significantly contributes to the understanding of the mechanism of action of PCMV/PRV on pig transplant survival. However, there are still open questions. It is still unclear, whether PCMV/PRV can infect baboon cells. Using PCR and immunohistochemical methods, PCMV/PRV was found in all organs of the transplanted baboon^27^, however it is unclear whether the cells stained positive with a specific antiserum against PCMV/PRV are disseminated PCMV/PRV-producing pig cells or infected baboon cells. It is also unclear whether PCMV/PRV can infect human cells. One report claimed infection of human cells^37^, whereas another report showed that human cells cannot be infected with PCMV/PRV^38^. Although not proven, reduction of the pig transplant survival may also happen in humans when PCMV/PRV-positive pig organs will be transplanted. Since different non-human primate species (cynomolgus monkeys, baboons) showed a similar reduction of the survival time (ref. 28, 29 and this study), it seems very likely that the same may happen in humans. There are other factors which may contribute to the reduction of pig transplant survival time. A consumptive coagulopathy was often observed in non-human primates with pig xenotransplants^34,39-41^. Although in vitro an activation of the porcine tissue factor (TF) in porcine aortic endothelial cells by a PCMV/PRV infection was observed, no correlation between TF expression and PCMV/PRV infection was observed in vivo^34^. An enhanced expression of ICAM-1 (intracellular adhesion molecule I) and MHC (major histocompatibility complex) class II in the pig transplant in baboons suggests an activation of endothelial cells^28^. PCMV/PRV is like many other viruses immunosuppressive and was shown to modulate the expression of immune-related genes in pig immune cells^22^. Furthermore, a PCMV/PRV infection of pigs is usually associated with opportunistic bacterial infections on the basis of the suppressed immune system^23^ and a transcriptome analysis of PCMV/PRV-infected thymuses showed an up- and downregulation of immune-regulatory genes^24^. However, considering the strong pharmaceutical immunosuppression of the animals in order to prevent rejection of the pig organ, the immunosuppressive properties of PCMV/PRV may not be important for the PCMV/PRV-induced pathogenesis.

The finding that IL-6 and TNF increased in the blood of baboons with PCMV/PRV-positive hearts is of great interest. IL-6 is one of the most important cytokines during an infection, along with IL-1 and TNF^42^. TNF is a cytokine involved in systemic inflammation and is a critical effector molecule in the immune response to viral pathogens^43^. TNF is able to inhibit viral replication and respond to sepsis via IL1- and IL6-producing cells.

Despite this seemingly overwhelming evidence for the role of PCMV/PRV, there may also be other factors contributing to the reduced survival rates. As explained elsewhere in detail^1^, xenotransplantation experiments in groups I and II failed because of insufficient organ preservation and overgrowth of the transplant, respectively. Baboons P and Q (group III) were euthanized because of beginning multi-organ failure, which might be attributed to other causes than PCMV/PRV: Baboon P had received a graft of borderline small size and baboon Q developed recurrent pericardial effusions. Relative size mismatch may lead to systolic heart failure, whereas recurrent effusions may lead to pericardial tamponade and diastolic insufficiency, both eventually causing multi-organ failure. Microbial translocation and opportunistic infections were observed in animals P and Q; both are common after infection with immunosuppressive viruses including human immunodeficiency virus (HIV)^44^, but can also be attributed to immunosuppressive treatment by itself.

It was clearly shown that PCMV/PRV replicates in the transplanted heart (Figure 1, Table 1). Already at day three an increase of the copy number in the heart was observed, reaching much higher copy numbers later. It may be suggested that the pharmaceutical immunosuppression given to prevent rejection and possibly the PCMV/PRV-induced immunosuppression plays an important role in allowing increased PCMV/PRV replication. For comparison, previous studies of the replication rate of HCMV in humans showed that its dynamics are rapid, with a doubling time of viraemia of approximately 1 day^45^. In analogy to the HCMV infection, where a clear threshold relationship was observed, e.g., the virus causes disease once a threshold value of viral load is exceeded^11^, such a threshold seems also important in the case of PCMV. In this context, a viral load of 2 to 3 million copies may be tolerable until day 30-40, whereas higher viral loads further reduce the survival time.

Since other viruses could also interfere with the survival time of the transplants, activation of BaCMV in the transplanted baboon and transmission of HEV, PERV, PCV1, PCV2, as well as PLHV1, 2 and 3 was analysed. All baboons used in these investigations carried BaCMV, which is common in baboons. In one case a strong activation of BaCMV after transplantation of a PCMV-positive pig heart was observed (baboon I)^27^. In all other cases a direct activation of BaCMV could not be observed. Concerning BaCMV activation, in another preclinical trial transplanting pig kidneys into baboons, it was also observed in control animals without transplantation, but with the corresponding immunosuppression, indicating that immunosuppression alone is able to activate BaCMV^46^.

HEV, genotype 3, is a well-known zoonotic virus, it is frequently transmitted to humans by undercooked meat or contact, but also by manure-contaminated fruits and water, and it induces chronic infection in immunosuppressed patients and severe liver disease in patients with an underlying liver failure^47^. However, until now no HEV transmission was reported in all preclinical and clinical xenotransplantation trials, including this study (Table 2).

PERV DNA was observed in the circulation of baboons following transplantation of a pig heart (Table 2). However, this observation appears to be due to persistent pig cell microchimerism.

PCV1 and PCV2 were not present in the donor pigs and, logically, could not be transmitted to the recipients. Meanwhile it was published, that four donor pigs (animals 5803, 5807, 6249, 6253) were infected with PCV3 and that this virus was transmitted in all four cases to the baboon recipients (baboons O, N, P, Q)^48^. PCV3 is a newly described member of the virus family *Circoviridae*, it is highly distributed among farms pigs and wild boars worldwide (for review see Reference 49). Similarly to the situation with PCV2, PCV3 was found in healthy animals as well as in animals suffering from different diseases, suggesting that coinfections with other viruses are necessary for the pathogenic potential. PCV3 does not seem to influence the survival time because baboon with the highest survival time (animal N - 182 days, 72 - O days) and animals with low survival times (P - 14 days, Q - 26 days) were infected with PCV3. In contrast, there was a clear correlation between the infection with PCMV and survival (14 and 26 days versus 182 and 195 days).

Although all donor pigs were infected with PLHV-1 and PLHV-2, no transmission to the baboon recipients was observed, PLHV-3 was not found in the donor pigs. PLHV-1, -2, and -3 belong to the subfamily Gammaherpesvirinae in the Herpesviridae family. The pathogenicity of PLHV in pigs under natural conditions is still unclear. Under experimental conditions PLHV-1 is associated with post-transplant lymphoproliferative disease (PTLD) in miniature pigs following allogeneic haematopoietic stem cell transplantation^50,51^. The clinical symptoms of experimental porcine PTLD, such as fever, lethargy, anorexia, high white blood cell count and palpable lymph nodes, are similar to those of human PTLD, a serious complication of solid organ and allogeneic bone marrow transplantation, which was linked to a human gammaherpesvirus, Epstein-Barr virus (human herpes virus-4, HHV-4)^52^.

In order to prevent transmission of porcine viruses after xenotransplantation, elimination programs had been proposed which are based on selection and isolation of virus-negative animals, vaccination or treatment with an effective antiviral drug (both are not available in the case of PCMV/PRV), early weaning, colostrum deprivation, Caesarean delivery or embryo transfer. Although PCMV/PRV can be transmitted via placenta^53,54^, successful elimination can be achieved by early weaning^35,55^, providing virus-free animals for a safe xenotransplantation.

## Methods

### Animals and transplantations

All details of the donor pigs, the baboon recipients and the transplantation procedures were described in reference 1.

### Immunosuppression

All baboon recipients received an immunosuppression including an induction therapy with an anti-CD20 antibody, anti-thymocyte-globulin and a monkey-specific anti-CD40 monoclonal antibody or humanized anti-CD40PASylated as described in detail^1^. The group III baboon recipients analysed here were weaned from cortisone at an early stage and received antihypertensive treatment. In addition, a temsirolimus medication was applied.

### Plasma and blood collection

Blood sampling from adult sows was performed without sedation under manual fixation. Whole blood was drawn from the jugular vein with single-use needles (Ehrhardt Medizinprodukte, Geislingen, Germany) into lithium heparin and serum Monovettes (Sarstedt, Nümbrecht, Germany). Blood from the baboons was taken using a central venous catheter.

### Ethics Statement

Both the generation of transgenic animals, as well as interventions on re-cloned animals, were performed with permission of the local regulatory authority. Applications were reviewed by the ethics committee according to §15 TSchG German Animal Welfare Act. The xenotransplantation experiment was approved by the Government of Upper Bavaria, Munich, Germany. Housing, feeding, environmental enrichment, and steps taken to minimise suffering, including the use of anaesthesia and method of sacrifice, was in accordance with the recommendations of the Weatherall report “The use of non-human primates in research”.

### Testing for PCMV/PRV

PCMV/PRV testing was performed as described^25^ using specific primers (Table 3). Briefly, DNA was extracted from sera, blood and organs of the pigs using the DNeasy Blood &Tissue kit (Qiagen GmbH, Hilden, Germany). DNA was quantified using a NanoDrop ND-1000 (Thermo Fisher Scientific Inc., Worcester, MA, USA). To screen for PCMV, a real-time PCR using described primers^26^ and the SensiFast probe no ROX kit was performed according to supplier recommendations (Bioline GmbH, Germany). DNA from formalin-fixed tissues was extracted by using the QIAamp DNA FFPE Tissue Kit (QIAGEN), following the manufacturer’s instructions. As the starting material was not paraffin-embedded but only formalin-fixed tissue, we started the process by cutting the tissue into small pieces, added buffer and proteinase and followed the manufacturer’s protocol by incubating at 56°C. A detection limit of 20 copies was determined for the reported PCR method^26^. Various amounts (25 - 250 ng) of DNA were used for testing. The reaction mixture contained 300 nM of both primers, and 250 nM of the probe (Table 3) in a final volume of 20 µL. The following conditions for amplification were used: denaturation at 95 C for 5 min, and 45 cycles of amplification with denaturation at 95 C for 15s, annealing at 56 C for 30s and extension at 72°C for 30s. Reporter fluorescence was measured using the CFX96 Toich Real-time PCR detection system (Bio-Rad, Hercules, CA, USA).

**Table 3.**
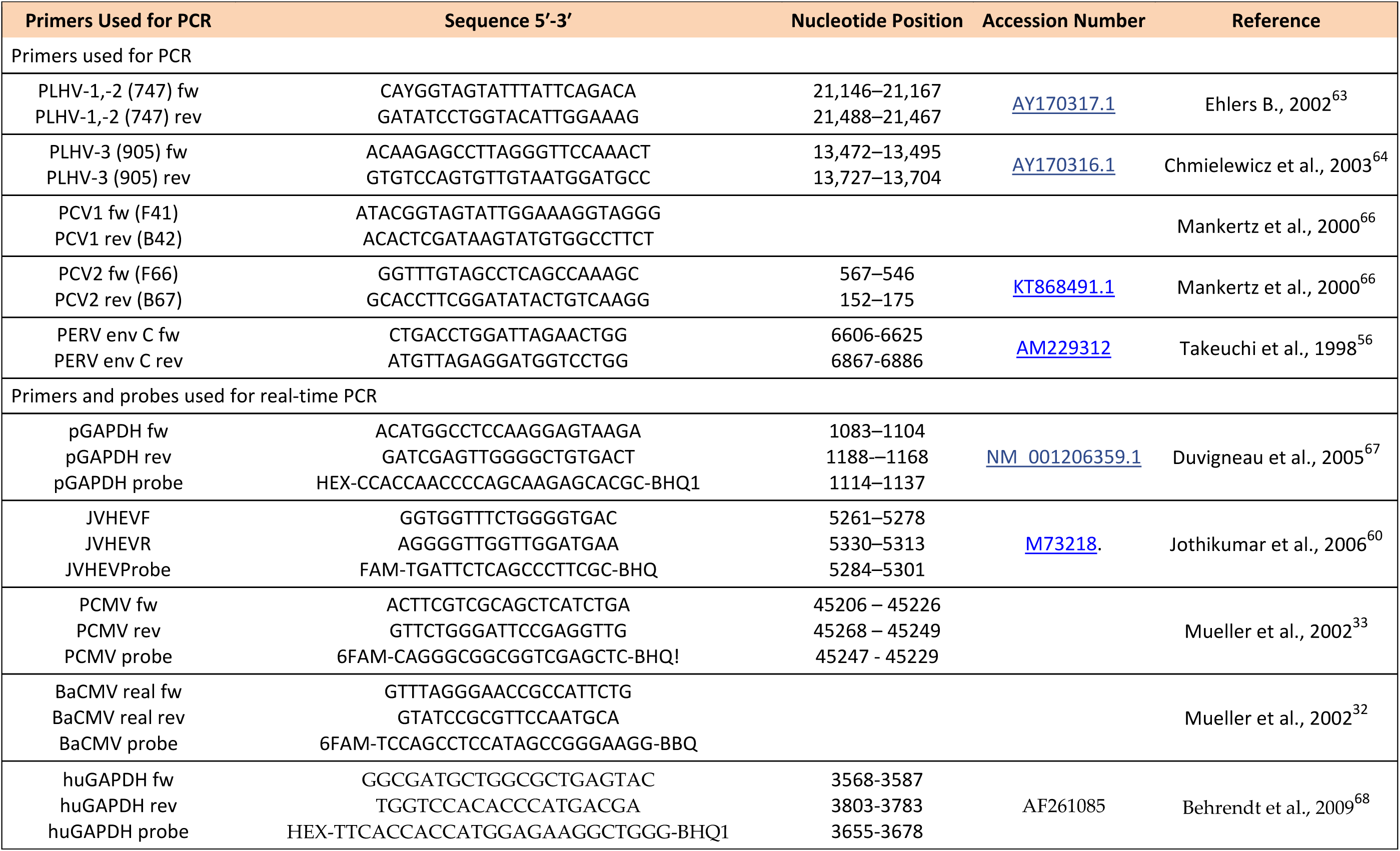
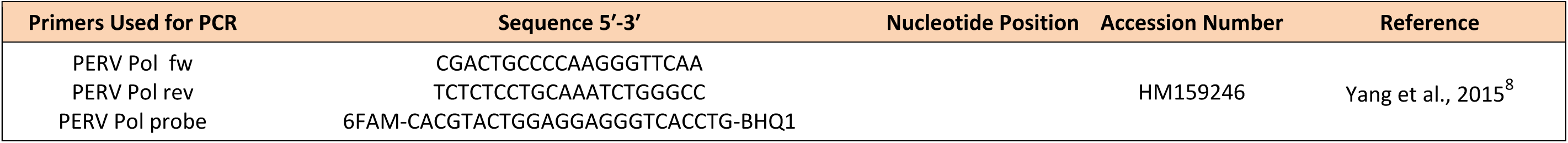
Primers and probes.

### Testing for BaCMV

To test for BaCMV, a real-time PCR was performed using described primers^32^ (Table 3), but the enzymes and conditions had been changed: Denaturation at 95°C for 5 min, and 45 cycles of amplification with denaturation at 95°C for 15s, annealing at 57°C for 30s and extension at 72°C for 30s.

### Testing for PERV

To test for PERV, a PCR was performed using primers for the pol sequence (Table 3) as described^4^. This PCR detects PERV-A, PERV-B and PERV-C, since the pol region is highly conserved. To test for PERV-C, a PCR was performed using primers specific for the env region of PERV-C (Table 3)^56,57^. A Western blot analysis was performed using recombinant p27Gag protein, p15E protein and gp70 protein as well as goat sera against these recombinant proteins as positive control as described^4,58^.

### Testing for HEV

To test for HEV, a real-time PCR was performed as described^59^ using specific primers^60^. Western blot analysis was performed using two overlapping recombinant ORF2 proteins of HEV genotype 3 as described^59^. On protein contained the immunodominant epitope (GT3-Ctr, aa 326-608, 32 kb)^61^, the other was with a glutathione-S-transferase tag (aa452-617, 44.5 kb, Orosperc, Ness Ziona, Israel).

### Testing for PLHV

To test for PLHV-1, PLHV-2 and PLHV-3 two PCR methods and specific primers (Table 3) were used as described^62-64^.

### Testing for PCV

To test for PCV1 and PCV2, a PCR method and specific primers (Table 3) were used as described^63^, using specific primers^65^.

### Measurement of cytokines

IL-6 release over time was studied in the Laboratoriumsmedizin of the Munich University using an ELISA (Roche, Elecsys IL-6).

The BD™ CBA Non-Human Primate (NHP) Th1/Th2 Cytokine Kit (BD Biosciences, Heidelberg) was used to measure IL-2, IL-4, IL-5, IL-6, TNF and IFN γ in baboon serum samples according to the manufacturer’s instructions. Serum samples were diluted 1:20 before use. Samples were acquired by use of a LSR II cytometer (BD Biosciences). To measure IL-10 serum levels the IL-10 Monkey ELISA Kit (Thermo Fisher, Waltham) was used according to the manufacturer’s instructions.

### Measurement of coagulation

The PAI-1/tPA complexes were measured in EDTA plasma samples by an in-house developed Bio-Plex assay using xMAP technology, as described previously^31^. Briefly, mouse anti–PAI-1 antibody was coupled to carboxylated polystyrene beads (Luminex) using a Bio-Plex amine coupling kit (Bio-Rad). The coupled beads were then incubated with samples and bound PAI-1/tPA complexes detected using biotinylated rabbit anti-tPA antibody (Molecular Innovations) and streptavidin-PE (Qiagen). Measurement and data analysis were performed with a Bio-Plex 100 array reader and the Bio-Plex Manager software version 6.1.

### Tissue processing and histological staining

All myocardial specimens were fixed in 4% neutral-buffered formalin and embedded in paraffin. Tissue sections were then stained with hematoxylin and eosin.

### Western blot analysis

To detect antibodies against PERV in baboons, recombinant surface envelope protein gp70, transmembrane envelope protein p15E and core protein p27 were used as described^4,58^. As control goat antisera against these recombinant proteins were used. Secondary anti-goat IgG and anti-human antibodies were used and the assay was developed using alkaline phosphatase. To detect antibodies against HEV in pigs and baboons, a recombinant genotype 3 (GT3) ORF 2 core antigen^61^ and a recombinant 44.5 kDa protein with a glutathione-S transferase (GST) tag fused to the ORF2 fragment (Prospec, Ness Ziona, Israel) was used^59^. Alkaline phosphatase conjugated antibodies were used and the reaction was developed using NBT (nitro-blue tetrazolium chloride)-BCIP (5-bromo-4-chloro-3’-indolyphosphate p-toluidine salt) substrate.

## Data availability

The data that support the findings reported herein are available on reasonable request from the corresponding author.

## Acknowledgments

This research was supported by the Deutsche Forschungsgemeinschaft, TRR127.

## Authors contribution

L.K. and U.F. performed the detection methods for all viruses, M.L., B.Re., M.M., J.R., T.M., P.B. and J-M.A. performed the transplantations, collected the samples for virus testing and arranged IL-6 testing. E.W. provided the multitransgenic pigs used for xenotransplantation. B.Ro. and C.S-H. performed the cytokine analyses, A.M., F.L., and R.R. performed the coagulation studies. C.W. performed the histological investigation, J.D. planned the virological part of the study and was writing the original draft preparation, L.K., U.F., M.L., B.Ro., J-M.A, A.M., F.L., N.S., R.R, C.W., and E.W. were reviewing and editing the manuscript, J.D., B.Re., R.R. and E.W. were supervising the project; J.D., R.R., P.B., B.Re. and E.W. were responsible for funding acquisition.

## Additional information

**Supplementary Figures** accompanies this paper at

## Competing interests

All authors declare no competing interests.

## Legends

**Supplementary Figure 1.**
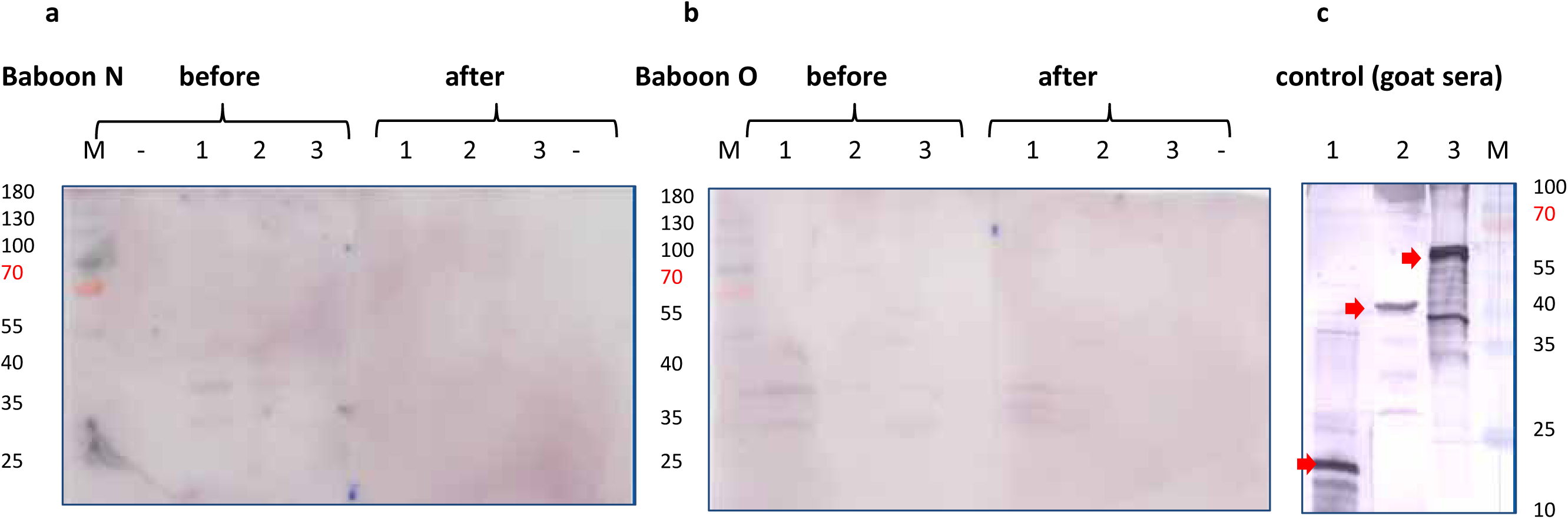
Western blot analysis of sera from baboon N and O before transplantation (before) and at the end of the experiment (after). **a** serum from baboon N, **b** serum from baboon O, **c** control sera obtained in goats against recombinant PERV antigens. 1, recombinant p15E, 2, recombinant p27Gag, 3, recombinant gp70. The red arrows indicate the recombinant proteins. The blots were overexposed to increase sensitivity and show unspecific bands.

